# Whole tissue spatial cellular analysis reveals increased macrophage infiltration in pancreata of autoantibody positive donors and patients with type 1 diabetes

**DOI:** 10.1101/2025.09.13.676053

**Authors:** Estefania Quesada-Masachs, Samuel Zilberman, Tiffany Chu, Sara McArdle, William B. Kiosses, Sakthi Rajendran, Mark A. Atkinson, Mehdi A. Benkahla, Zbigniew Mikulski, Matthias von Herrath

**Affiliations:** Diabetes Research Institute, Leonard M. Miller School of Medicine, University of Miami, Miami, FL; La Jolla Institute for Immunology, La Jolla, CA, 92037 USA; Diabetes Institute, University of Florida, Gainesville, FL, 32608 USA

## Abstract

While extensive efforts have characterized lymphoid populations that contribute to pancreatic ‘insulitis’ in type 1 diabetes, significant gaps remain regarding key properties of macrophages in this inflammatory lesion. Hence, we quantified the spatial distribution of macrophages infiltrating human pancreatic tissue sections across disease states. Whole slide images of immunofluoresent stainings were analyzed with a semiautomated machine learning approach to study the distribution of macrophages throughout pancreatic sections from non-diabetic, auto-antibody positive (Aab+), and type 1 diabetes organ donors. Across the entire tissue section, macrophage infiltration was highest in donors with type 1 diabetes, and appeared contingent on the prescence of insulin-containing islets. Aab+ donors had higher macrophage densities in islet and peri-islet regions than non-diabetic individuals, but there was large case-by-case variation. Additionally, a greater proportion of macrophages highly expressed HLA class II (HLA-II) near and within islet regions of the type 1 diabetes group, which correlated with the proportion of stressed beta cells expressing HLA-II. The higher density of macrophages in the Aab+ and type 1 diabetes donors, taken together with higher expression of HLA-II in the type 1 diabetes pancreas, suggests that macrophages play a pivotal role in human type 1 diabetes initiation and progression.

**Article Highlights:**

- Macrophage infiltration is increased in all pancreatic regions of patients with type 1 diabetes, and this infiltration appears contingent on the presence of insulin-containing islets.
- Aab+ donors have higher macrophage densities in islet and peri-islet regions than non-diabetic individuals, but there is large case-by-case variation within the Aab+ group.
- The proportion of macrophages highly expressing HLA-II is increased near and within islets only in patients with type 1 diabetes, and this correlates with the proportion of stressed beta cells expressing HLA-II within the islet.
- Macrophages are linked to human type 1 diabetes initiation and progression, hence forming a rationale to examine therapeutic strategies involving this cellular population.

---

Type 1 diabetes is an organ-specific autoimmune disease characterized by the destruction of insulin-producing pancreatic beta cells. Currently, it is commonly accepted that, in patients with type 1 diabetes, infiltrating autoreactive CD8+ T cells are the primary mediator of beta cell destruction [1–3]. However, there has been an increased focus on understanding the involvement of the innate immune system—especially macrophages—in disease initiation and progression [4].

Macrophages function as professional antigen presenting cells primarily through the expression of HLA class II (HLA-II). As a broad classification, activated macrophages are characterized as having either an M1 (proinflammatory) or M2 (immunoregulatory) phenotype but their high plasticity allows many intermediate functional states [5]. It has become evident that macrophage phenotype and function are dynamic and quickly respond to changing tissue and disease-specific factors [6]. As such, in an organ-specific disease like type 1 diabetes it is particularly relevant to study macrophages *in situ*.

While resident macrophages are present in islets of non-diabetic individuals [7, 8] and have been found to be important for islet development and homeostasis in healthy C57BL/6 mice [9, 10], studies in non-obese diabetic (NOD) mice suggest that macrophages may be a key mediator of the early stages of disease progression—potentially even before diagnosis or onset of symptoms [11]. Patients who are seropositive for multiple or even a single autoantibody (Aab+) are known to be associated with an increased risk of developing type 1 diabetes [12]. Studying pancreatic sections of donors with autoantibodies is especially important for understanding macrophage population dynamics throughout early disease stages in human type 1 diabetes. In the present study we analyzed, characterized, and compared macrophage populations and their spatial distribution throughout pancreatic sections from type 1 diabetes, Aab+, and non-diabetic organ donors.

## Research Design and Methods

### Pancreatic Organ Donors

Formalin-fixed paraffin-embedded (FFPE) tissue sections of human pancreas were obtained through the Network of Pancreatic Organ Donors with Diabetes (nPOD) program, with informed consent from donor families obtained by an Organ Recovery Organization partner of that organization (https://www.jdrfnpod.org/for-partners/organ-recovery-partners/). Pancreatic nPOD sections were from 15 brain-dead organ donors (non-diabetic (n=5), Aab+ (n=5) and type 1 diabetes donors (n=5)) matched for age, sex, and body mass index (demographic and clinical information in Supplementary Table 1).

### Immunofluorescence Staining

Immunofluorescence staining was performed following procedures outlined in our prior study [13]. In summary, FFPE pancreatic sections were deparaffinized with two baths of pro-par clearant for 10 minutes each and then rehydrated to water in descending concentrations of ethanol (100%, 90%, 70%, 50%, 0%) for 5 minutes each. Antigen retrieval was performed by incubating slides in citrate buffer (pH 6.0) at 95°C for 20 minutes. Slides were positioned in a 3D printed immunostaining slide rack (https://3dprint.nih.gov/discover/3dpx-012172) for the staining procedure. Tissues were blocked with 10% goat serum for 1 hour. Next, slides were incubated overnight with primary antibody mouse anti-human HLA-II 1:100 (DP, DQ, DR; DAKO) at 4°C, and the next morning with secondary antibody goat anti-mouse AF647 (Jackson) for 1 hour at RT. Slides were then incubated overnight with rabbit anti-CD68 monoclonal 1:200 (Abcam) at 4°C, and subsequently labelled with secondary antibody goat anti-rabbit AF555 (Invitrogen) for 1 hour at RT. After that, slides were incubated with mouse anti-insulin-AF488 1:400 (eBioscience) for 1 hour at RT. Finally, slides were counterstained with Hoechst 33342 (Life Technologies) for 10 minutes, and then coverslips were mounted using Pro-long gold anti-fade. The full panel of antibodies and staining reagents used can be found in the key resources table.

### Image Acquisition

Images of whole pancreatic tissue sections were obtained using a Zeiss AxioScan.Z1 widefield slide scanner equipped with a 20x (0.8 NA) objective (Z-stacks were acquired with a 0.5 μm step size). Additionally, high-resolution images were obtained of randomly selected islets from each case across the tissue section and were acquired with a Zeiss laser scanning confocal microscope LSM880 with an oil 40x (1.4 NA) objective using a 0.3 μm step size (8 z-steps per imaging field). All 8-bit images were acquired using the full dynamic intensity range (0-256).

### Image Analysis

The widefield whole slide image Z stacks were flattened to a single best-focused plane using a custom Zen macro previously described [13]. The processed images were imported into QuPath [14]. Pixel size was homogenized according to original data (0.325 µm), channel names were specified, minimum intensity thresholds for each channel were calculated. We optimized the built-in automatic cell detection based on the Hoechst (nuclei) signal and trained machine learning object classifiers (random trees) for each signal (insulin, HLA-II, and CD68). After composite classifiers were generated to detect double and triple positive cells (CD68 + HLA-II and insulin + HLA-II), a second Groovy script was programmed to retrieve the total number of cells and the number of cells expressing each marker throughout the entirety of every tissue section. Macrophage density maps were created, overlaid on the original tissue images using the built-in tool in QuPath and were used to visually identify macrophage “hotspots”.

For the regional analysis, we trained a pixel classifier (random trees) to automatically identify the ICI regions based on the insulin signal, and then manually corrected any mistakes, and IDIs were manually identified in each case by selecting for the islet-shaped regions characterized by lower autofluorescence in the background Hoescht signal. Additionally, 15 exocrine regions were randomly selected across each tissue section. Next, we programmed a Groovy script to automatically draw a peri-islet region of 10 µm around the perimeter of each islet region. For each region of interest (islet, peri-islet, and exocrine), the area, the total number of cells, and the number of cells expressing each marker were calculated. For the expanded islet ring analysis, we wrote a custom script to automatically draw eight concentric rings with a 25 µm radius expanded from the islet boundaries. Within each ring and the islet regions themselves, the area, the total number of cells, and the number of cells expressing each marker were calculated. In total, 7415 islets with their respective peri-islet or expanded ring regions were analyzed and, from the entire tissue section, the data of more than 24 million cells (>1 million macrophages (CD68+ cells)) was retrieved.

### Statistical Analysis

Prism 9 software (GraphPad) was used for statistical analysis. We performed a descriptive statistical analysis first to determine mean, SD, median, mode, min-max values, fences, IQR and quartiles. Shapiro-Wilk tests were used to determine if the data was normally distributed. For the normally distributed data, we used ordinary one-way ANOVA to calculate p-values and identify significant differences between more than two groups. For non-normally distributed data, the non-parametric Kruskal-Wallis test was used to calculate p-values and identify significant differences between more than two groups. We also performed the Tukey’s test (for normally distributed data) or Dunn’s test (for non-parametric data) to assess multiple comparisons and specific differences between groups, when appropriate. Correlation analysis was performed using Spearman’s correlation test. P-values < 0.05 were considered significant and adjusted when applying multiple comparisons. Information about the specific statistical test used can be found in the figure caption corresponding to each figure.

## Results

### Macrophage infiltration is significantly increased in patients with type 1 diabetes

Cells visually identified as macrophages could be observed in all pancreatic sections from non-diabetic, Aab+, and type 1 diabetes donors. After segmenting cells and training an object classifier to identify CD68+ macrophages, we analyzed all cells in the 15 tissue sections (24,473,800 total cells were included in the analysis). Heat maps displaying the relative density of macrophages across the tissue (density maps) revealed increased macrophage infiltration in most type 1 diabetes and some Aab+ donors compared to the non-diabetic ones (Fig. 1A). Cell quantification of macrophages throughout the entire tissue section, noted that patients with type 1 diabetes had a statistically significantly increased percentage of total cells expressing CD68 compared to the non-diabetic and Aab+ patients. The mean percentage of cells expressing CD68 in the type 1 diabetes group was 8.6±4.7% compared to 3.2±2.7% and 1.5±0.5% in the Aab+ and non-diabetic groups, respectively (p=0.01; Fig. 1B). Concordantly, the density of CD68+ cells was higher in the type 1 diabetes group (856.3±508.2 cells/mm^2^) than in the Aab+ (312.8±268.4 cells/mm^2^) or non-diabetic (136.6±56.3 cells/mm^2^) groups (p=0.01; Fig. 1C). Notably, there was considerable variation in macrophage infiltration between cases within the type 1 diabetes and Aab+ groups. Variation in the type 1 diabetes group is attributed to an outlier (nPOD #6084), which, unlike sections from the other pancreatic donors in this group, contained zero insulin-containing islets (ICI). The Aab+ group also had notable variation in macrophage infiltration between cases. It appeared that nPOD #6123, #6197, and #6301 exhibited an increase in the pancreatic macrophage infiltration, closer to that of the type 1 diabetes group, while nPOD #6147 and #6167 were more similar to the non-diabetic group (Fig. 1A). Overall, case-wise deviations within each group can be noted from the full-tissue analysis (Fig. 1).

**Figure 1.**
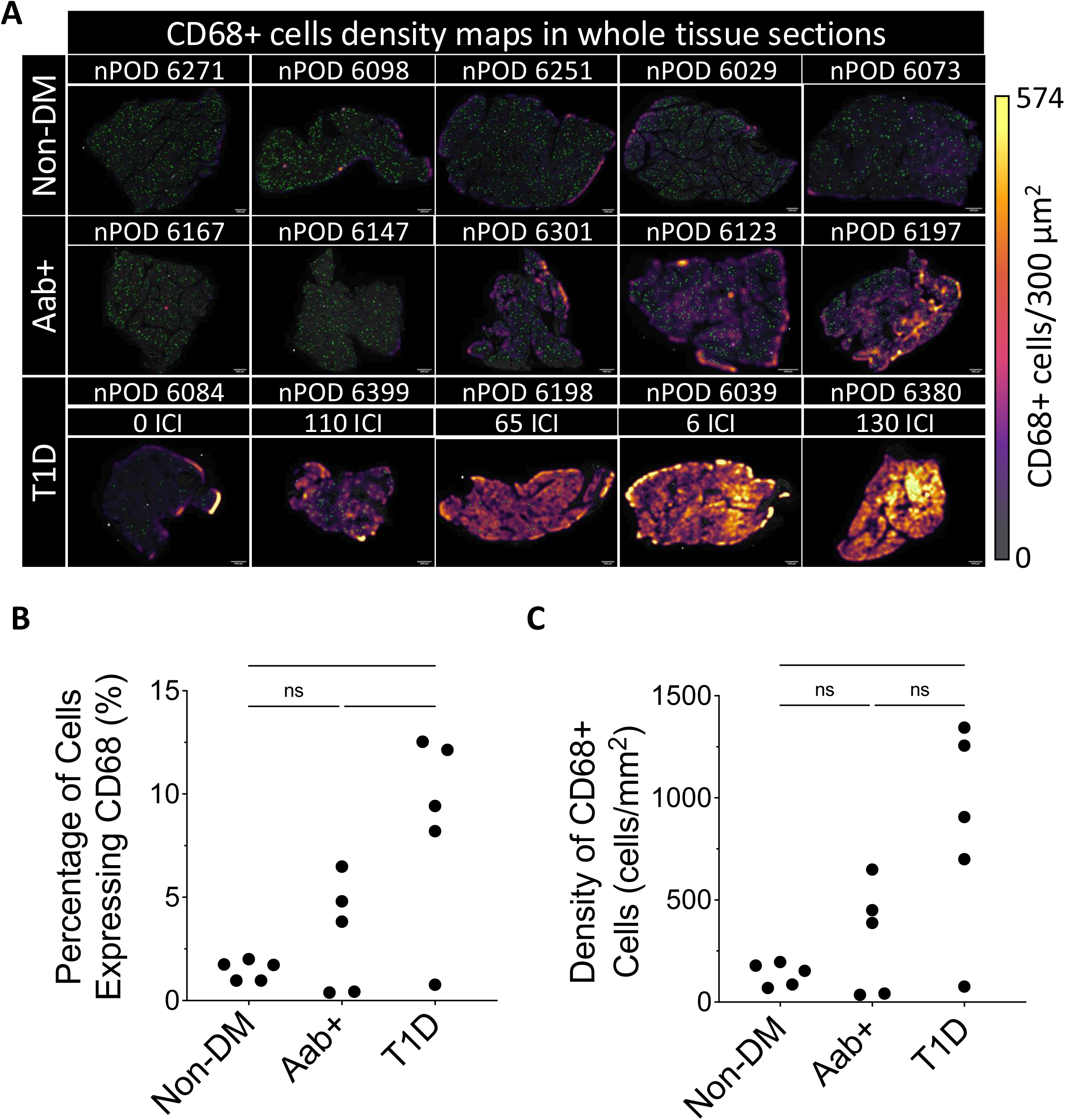
Macrophage infiltration is increased in the pancreata of donors with type 1 diabetes: **(A)** Images of each whole pancreatic tissue section with islet annotations highlighted in green, and heatmaps overlaid displaying macrophage density (scale bar, 2000µm). For each group, the cases are arranged from lowest to highest density (left to right). For each type 1 diabetes (T1D) case, the number of insulin-containing islets (ICI) are listed. Macrophage infiltration throughout the whole tissue is statistically significantly increased in patients with T1D as shown by **(B)** the percentage of total cells expressing CD68 (p=0.0101) and by **(C)** the density of CD68+ cells (p=0.0129). Each data point represents one case, and the mean + SD is shown. A total of 24,473,800 cells and 1,015,701 macrophages (CD68+ cells) were included in the analysis. Statistical significance was calculated using one-way ANOVA, and Tukey’s test for multiple comparisons (* p<0.03).

### Spatial distribution of macrophages in human pancreata

Macrophage distribution was first examined by analysing regions of interest: islet, peri-islet (10 µm radius), and selected exocrine regions (Supp. Fig. 1A). Quantification of the percentage of cells expressing CD68 indicated that macrophage infiltration was highest in the peri-islet regions regardless of disease status (for each disease group p<0.0001; Fig. 2A). CD68+ cells represented 19.6±14.3%, 9.5±8.1%, 3.9±6.9%, and 1.9±4.5% of the total cells in the peri-islet regions of the type 1 diabetes ICI, type 1 diabetes insulin-deficient islets (IDI), Aab+, and non-diabetic groups, respectively (Fig. 2A). Overall, the differences in the proportion of CD68+ cells between regions were less pronounced between the Aab+ and non-diabetic donors. In type 1 diabetes, ICIs exhibited a greater proportion of CD68+ cells than their exocrine regions (10.8±10.2% vs 7.9±4.7%) while IDIs had a lower proportion of CD68+ cells (1.8±2.5%). The islet regions of Aab+ donors also exhibited a greater proportion of CD68+ cells than the exocrine regions (2.5±7.5% vs 2.3±2.6%) while islets from the non-diabetic group had a lower proportion of CD68+ cells than the exocrine regions of this group (0.6±2.5% vs 1.0±0.7%). Overall, the type 1 diabetes group had the highest percentage of cells expressing CD68+ throughout all the tissue regions (exocrine, peri-islet, and ICIs) when compared to all other groups (Supp. Fig. 2). The only exception were type 1 diabetes IDIs, with a lower proportion of CD68+ cells than islets of Aab+ donors (1.8±2.5% vs 2.5±7.5%, respectively).

**Figure 2.**
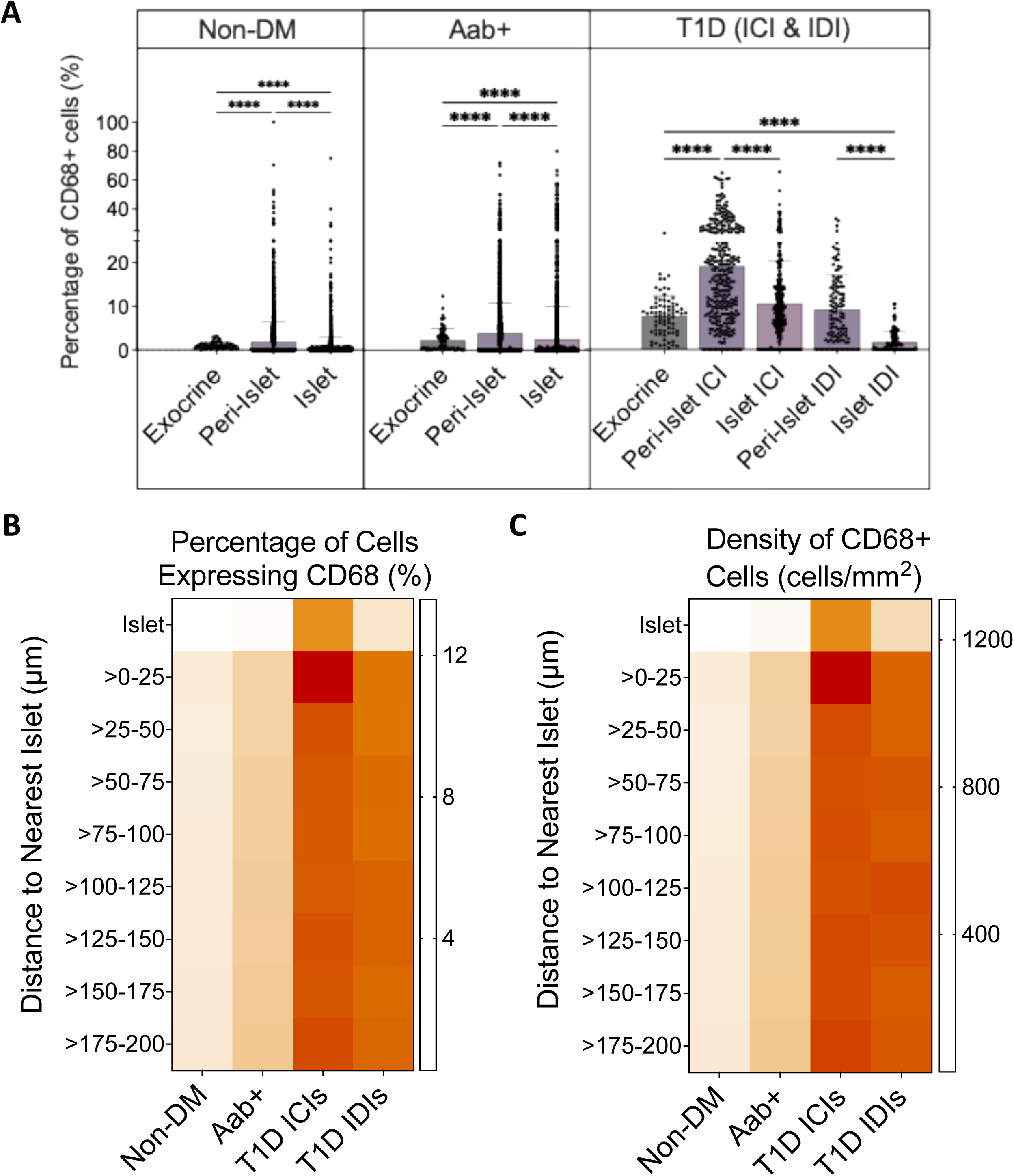
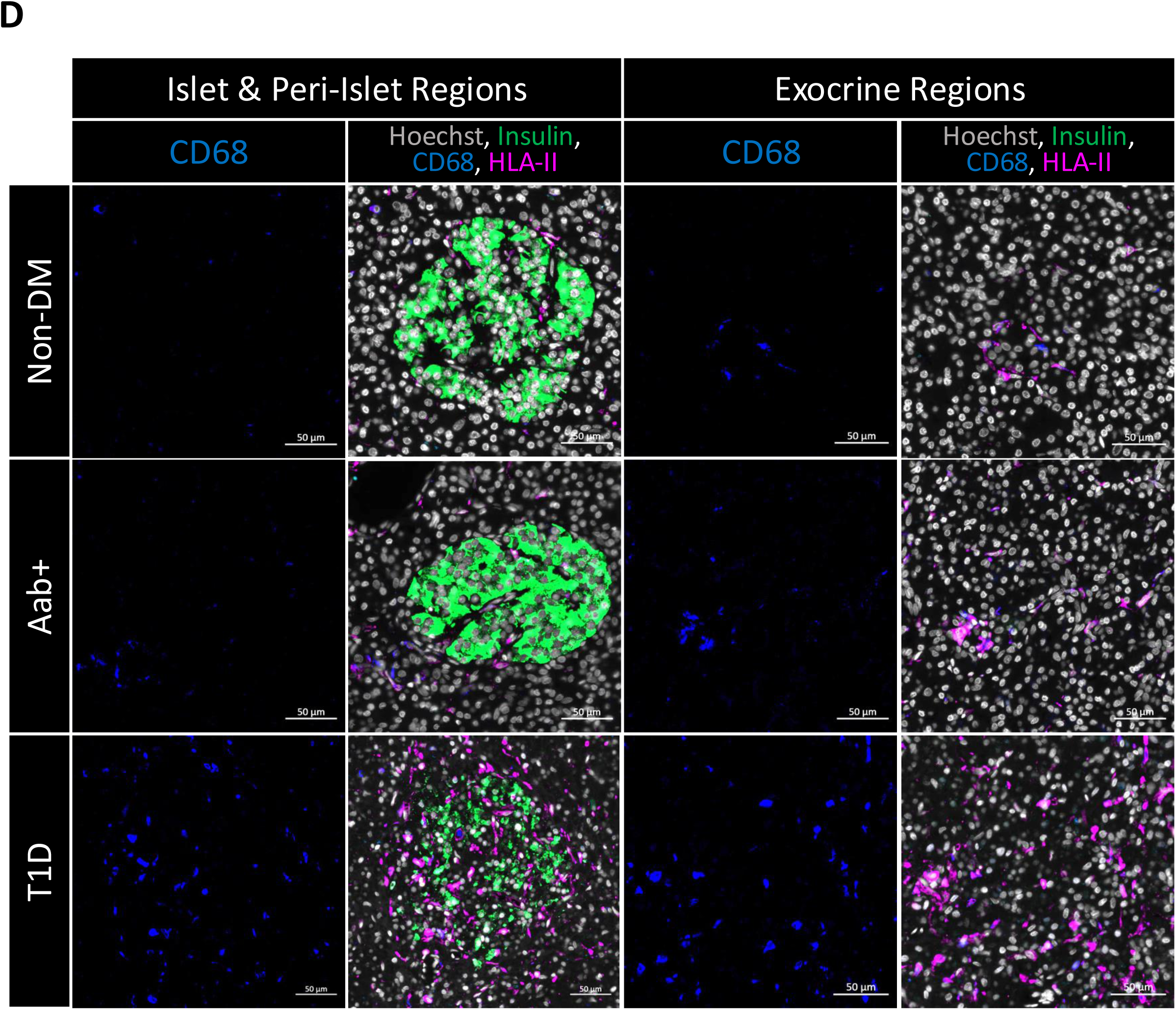
Spatial distribution of macrophages in human pancreata: Upon first spatial analysis using islet, peri-islet (10 µm radius), and selected exocrine regions, the **(A)** percentage of macrophages was statistically significantly highest in peri-islet regions regardless of disease status (Kruskal-Wallis p<0.0001 for non-diabetic, Aab+ and type 1 diabetes (T1D) ICIs and IDIs, multiple comparisons using Dunn’s test **** p<0.0001). Bars show mean + SD. Each data point represents one islet or region of interest. Further analysis of expanded islet regions made up of 8 concentric rings (each with a 25 µm radius) revealed that while **(B)** the percentage and **(C)** density of macrophage infiltration was higher around the islet than within it, this increase was also present beyond the immediate islet regions in all groups. Statistical significance was calculated using Two-way ANOVA test with p<0.0001 for both B and C. Multiple comparisons were assessed using Tukey’s test (B, C). **(D)** Representative images of the pancreatic regions (islet, peri-islet, and exocrine) from each donor group. Immunofluorescent antibody staining with hoechst –grey; insulin – green; CD68 – blue; HLA class II – magenta. Images were acquired with Zeiss Axioscan.Z1 slidescanner (20x objective).

To determine whether the localization of macrophages in the peri-islet regions extended beyond 10 µm creating a gradient of macrophage infiltration around the islet, 8 concentric ring regions (each with a 25 µm radius) reaching up to 200 µm away from every islet region were created (Supp. Fig. 1B). We observed statistically significant differences in the density and proportion of CD68+ cells between disease status (p<0.0001, Fig. 2B and 2C). Consistent with the findings from the first regional analysis, the percentage and density of CD68+ cells were higher in the first ring (within 25 µm of the islet perimeter) than within the islet region for all groups. Within the first ring, the proportion of CD68+ cells were 13.6±5.1%, 7.5±4.0%, 2.6±2.4%, and 1.3±0.4% compared with 6.2±4.2%, 1.6±1.5%, 0.4±0.5%, and 0.3±0.1% of the islet region for the type 1 diabetes ICI, type 1 diabetes IDI, Aab+, and non-diabetic groups, respectively (Fig. 2B). Likewise, comparison of the density of CD68+ cells between the islet region and the first ring found similar results with higher CD68 densities for the type 1 diabetes donors (Fig. 2C). The proportion and density of CD68+ cells were sustained at similar levels to that found in the first ring for the subsequent 7 rings for all groups except around the ICIs of the type 1 diabetes group. For the type 1 diabetes ICIs, after the initial spike in the proportion and density of macrophages within 25 µm of the islet perimeter, macrophage infiltration declined and stabilized in the subsequent 7 rings, but still remained elevated compared to that of the islet region. On a case-wise basis, the distribution of macrophages varied between donors of the same group (Supp Fig. 3A, B). Despite the case-wise variations, once all regions from each case are combined into their respective group, the percentage and density of macrophages in each ring region are approximately uniformly distributed (Supp. Fig. 4A, B).

**Figure 3.**
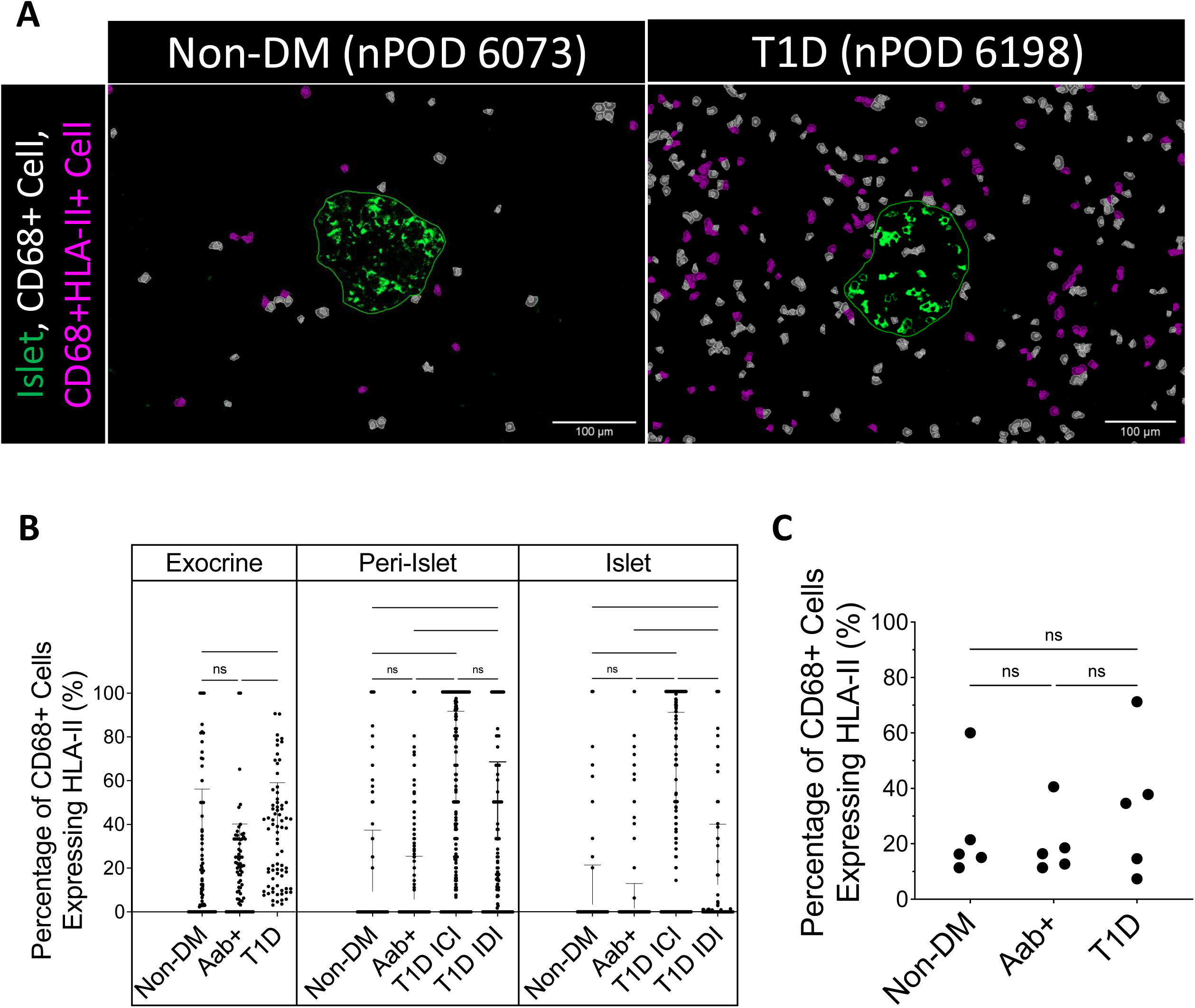
HLA-II is significantly upregulated by macrophages in and around islets of patients with type 1 diabetes: **(A)** Representative images of macrophage infiltration around islets (scale bar, 100µm), with insulin original signal (green), islet annotation (green outline), cell detections of macrophages CD68+HLA-II-(white), and CD68+HLA-II+ (magenta). **(B)** The percentage of macrophages expressing HLA-II (from the total of CD68+ cells) is increased in exocrine (p<0.0001), islet (p<0.0001) and peri-islet (10 µm radius) regions (p<0.0001) of patients with type 1 diabetes (T1D). Each data point represents one islet, peri-islet or exocrine region. Mean + SD is shown. Notably, this trend was not observed when considering **(C)** the percentage of macrophages expressing HLA-II across the whole tissue (p=0.5773). Each data point represents one donor. Mean + SD is shown. Statistical significance was calculated with Kruskal-Wallis (B) or One-way ANOVA (C), and Dunn’s test (B) or Tukey’s test (C) for multiple comparisons (*p<0.03, **p<0.002, ***p<0.0002 ****p<0.0001).

Representative images from islet, peri-islet, and exocrine regions of every group are consistent with these findings and demonstrate that macrophage infiltration is highest in the type 1 diabetes group throughout all regions, while that in Aab+ and non-diabetic cases appears to be more similar (Fig. 2D).

### HLA-II is significantly upregulated by macrophages located in and around islets of patients with type 1 diabetes

Our cell classification process allows for identification of cells co-positive for two or more markers. While type 1 diabetes cases appeared to contain more macrophages, it also appeared that a higher number of those macrophages expressed HLA-II (Fig. 3A).

Analysing the data from the exocrine, peri-islet, and islet regions, a much greater proportion of the macrophages highly expressed HLA-II in type 1 diabetes donors compared to the other groups. In the peri-islet regions, 49.2±42.0% and 32.6±35.7% of macrophages highly expressed HLA-II for the type 1 diabetes ICIs and IDIs, respectively, while only 5.7±19.6% and 9.3±27.9% of macrophages highly expressed HLA-II in peri-islet regions of Aab+ and non-diabetic groups, respectively (p<0.0001; Fig. 3B). A similar trend was observed in the islet regions where the percentage of macrophages highly expressing HLA-II was greater for the type 1 diabetes ICIs and IDIs than the Aab+ and non-diabetic islets (44.9±45.7% and 12.3±27.5% vs 1.6±11.3% and 3.4±17.8% respectively) (p<0.0001; Fig. 3B).

While there was a greater proportion of macrophages highly expressing HLA-II in the selected exocrine regions of the type 1 diabetes group (35.0±24.1%) compared to either the Aab+ (20.1±20.1%) or non-diabetic (25.1±31.0%) groups (p<0.0001; Fig. 3B), these differences were less pronounced. This translates to the finding that no relevant differences exist between groups in the percentage of macrophages highly expressing HLA-II when considering the entire pancreatic tissue section of each case, although the overarching trend remains. Throughout the whole tissue, 33.1±24.9%, 19.9±11.9%, and 24.8±20.0% of macrophages highly expressed HLA-II in the type 1 diabetes, Aab+, and non-diabetic groups, respectively (p=0.577; Fig. 3C).

### Islet and peri-islet infiltration of HLA-II+ macrophages correlate with beta cell expression of HLA class II in type 1 diabetes islets

Considering the high proportion of macrophages in and near the islets of type 1 diabetes cases, the impact these cells might have on the beta cells was investigated in the type 1 diabetes donors. Macrophages could be frequently observed in close-contact with the insulin-producing beta cells of the islet—some of which highly expressed HLA-II (Fig. 4A). There was a weak correlation of the percentage of cells expressing CD68 in islet and peri-islet regions with the percentage of beta cells which highly expressed HLA-II (r=0.28; 95% CI [0.17, 0.38]; p<0.0001; Fig. 4B). However, there was a strong correlation of the percentage of cells co-expressing CD68 & HLA-II in the islet and peri-islet regions with the percentage of islet beta cells which expressed HLA-II (r=0.74; 95% CI [0.68, 0.78]; p<0.0001; Fig. 4C).

**Figure 4.**
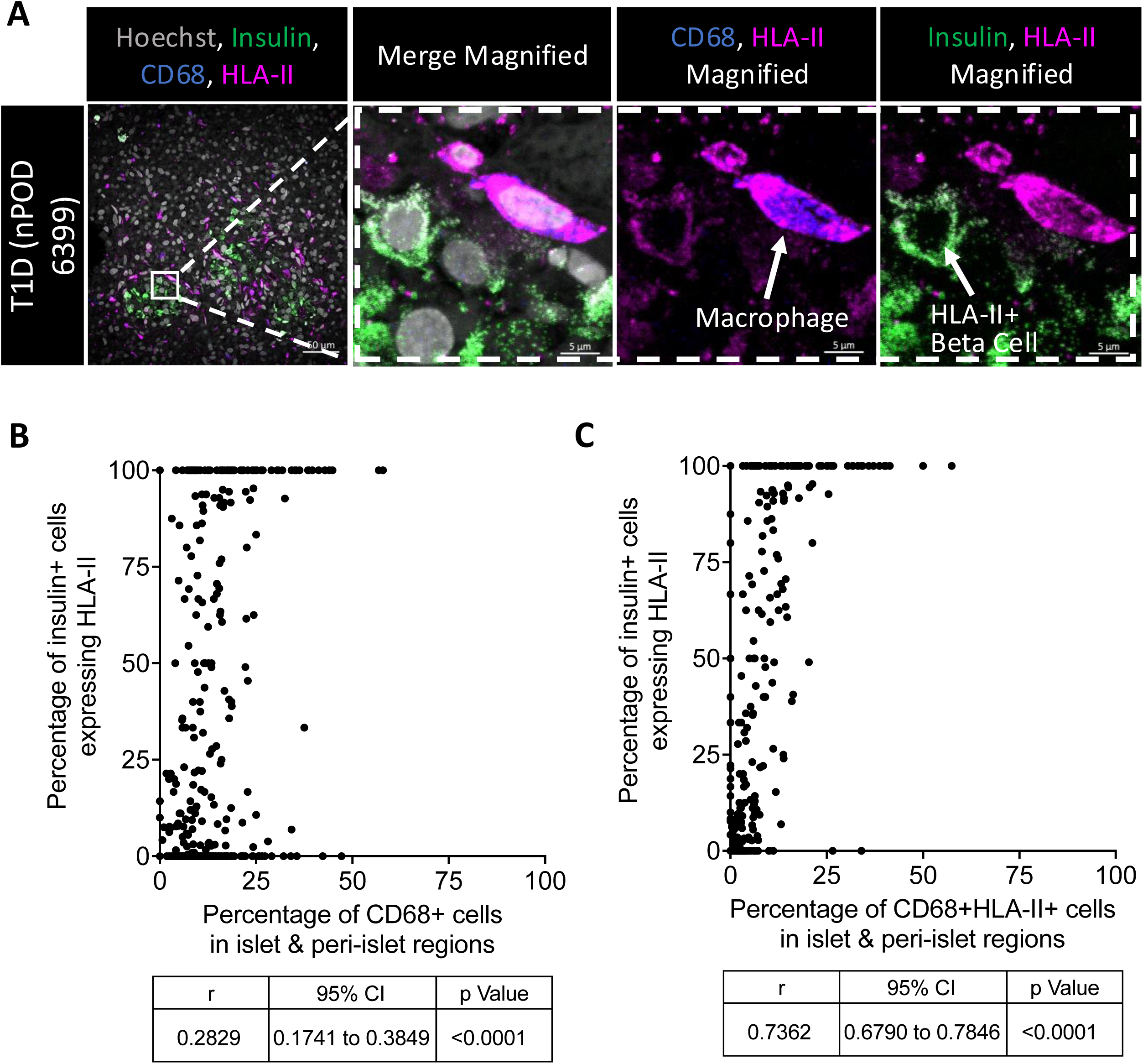
Islet and peri-islet infiltration of HLA-II+ macrophages correlates with beta cell expression of HLA class II in the T1D pancreata: **(A)** Representative high-resolution confocal image (scalebars, 50µm and 5µm) of a macrophage (CD68+) in close proximity to a beta cell (insulin+) expressing HLA class II (hoechst – gray; insulin – green; CD68 – blue; HLA class II – magenta). **(B)** Correlation of the percentage of CD68+ cells (macrophages) in islet and peri-islet regions with the percentage of beta cells expressing HLA-II in islets of type 1 diabetes (T1D) donors. **(C)** Correlation of the percentage of cells that co-express CD68 and HLA-II (activated macrophages) in islet and peri-islet regions with the percentage of beta cells expressing HLA-II in islets of T1D donors. Correlation coefficient, 95% confidence interval, and statistical significance were calculated with Spearman’s nonparametric correlation test.

## Discussion

Past studies have observed that macrophages are consistently one of the largest populations of immune cell in the pancreas of patients with type 1 diabetes [8, 17, 18]. Our finding that macrophage infiltration is significantly increased in the pancreas of patients with type 1 diabetes complements those previous findings and demonstrates that high levels of macrophages distinguish pancreata of patients with type 1 diabetes from that of non-diabetic individuals. While CD68 is not a definitive macrophage marker—indeed other monocytes, dendritic cells, and APCs are known to express CD68—the protein is commonly expressed in macrophages across activitation states for its critical role in phagocytosis, and has been previously used as a pan-macrophage marker in many human tissues, including the pancreas, across various disease contexts [15–18]. These macrophages may be originating from proliferating tissue resident macrophages, or they may be recruited from circulating monocytes [19, 20].

We observed increased macrophage infiltration across all type 1 diabetes pancreatic sections except that of nPOD #6084, which exhibited macrophage levels more similar to the non-diabetic group. The only differentiating factor of this case from the other type 1 diabetes cases was that it did not contain any ICIs, and this suggests that the increased presence of macrophages might be dependent on the presence of insulin. It is plausible that once all functional beta cell mass is lost, the lack of autoantigenic targets will dissipate inflammatory stimuli and infiltrating immune cells might recede from the insulin deficient regions. Even though more research is necessary to validate this notion, other groups previously described a significant decline in immune infiltrates of IDIs when compared with ICIs in type 1 diabetes [8, 21].

Multiple studies in mice demonstrated that islet macrophages are in intimate contact with beta cells and suggest that they could be key mediators of the early stages of disease progression [22, 23]. Resident macrophages are in proximity with the islet vasculature and beta cells, and have been observed to engulf beta cell secreted vesicles containing insulin and insulin catabolites, in both B6 and NOD mice [24]. In our study, we observed higher proportions of macrophages located in close proximity to the islets regardless of the disease status. This reinforces previous data suggesting that macrophages participate in the homeostasis of the islets [25, 26]. We observed a lower proportion of macrophages inside the islet when compared to the proportion of them around the islet. One potential explanation for this finding is that the peri-islet basal membrane creates a barrier that limits the immune infiltration into the islet [27]. Some evidence suggests that the localization close to the islets of a macrophage subpopulation that secrete proteases correlates with sites of loss of peri-islet basal membrane in type 1 diabetes; this membrane loss seems to be a critical step for leukocyte penetration into the islet, and hence for early type 1 diabetes progression [28].

Selective depletion of islet resident macrophages in NOD mice at three weeks-of-age resulted in diminished early infiltration of CD4+ T cells to the islets and reduced the incidence of autoimmune diabetes [11], highlighting the significance of resident macrophages in disease initiation. In mice islet and peri-islet infiltration with macrophages preceded the disease onset [29]. The elevated level of macrophages we observed in some of the Aab+ donors compared to the non-diabetic group, evidences this idea that macrophages might be involved in initiating the autoimmune attack in the human pancreas, and that they are present even before diagnosis or overt symptoms of type 1 diabetes. The high level of variation in macrophage infiltration within individuals in the Aab+ group could not be correlated with any reported clinical characteristic of the patients (data not shown) but could point to the incomplete penetrance of type 1 diabetes among patients with autoantibodies or the differing timelines in which Aab+ individuals begin to lose beta cell mass [12]. Spatial distribution of macrophages was heterogeneous between and within cases of every group, which has previously been reported in type 1 diabetes pathology based studies [30]. Heterogeneity is likely attributable to individual differences in the disease evolution, and to the lobularity in the distribution of the autoimmune attack within the pancreas [31].

While our study is limited by the relatively low number of samples we have analyzed (n=15; 5 per group), our semi-automated machine learning approach to quantifying the spatial distribution of macrophages throughout entire pancreatic tissue sections represents a significant advancement in *in situ* image analysis methodology. We can provide an extraordinary level of detail into the cellular microenvironment of a tissue or within regions of interest inside the tissue (e.g.., islets). Another study using imaging mass cytometry found that macrophages were the most common immune cell in islet and peri-islet regions of non-diabetic and type 1 diabetes donors [7]. Our current analysis confirms and supports the claim that macrophages are primarily positioned within and around the ICIs of the type 1 diabetes pancreas. However, type 1 diabetes donors also had higher proportions of macrophages in the selected exocrine regions when compared with the other two groups. This increased infiltration of immune cells in the exocrine tissue of patients with type 1 diabetes has been previously described [32]. Clearly, immune infiltration is not confined to just the islets in type 1 diabetes: the exocrine tissue also experiences changes in the local microenvironment. For the non-diabetic and Aab+ groups, while there are macrophages present in and near islets, the differences between the relative number of islet-infiltrating macrophages and exocrine macrophages are not as distinctive as that of the type 1 diabetes group.

The prevailing dogma of the past decades has proposed that type 1 diabetes is a disease of the adaptive immune system. However, the limited efficacy of preventative immunosuppressive therapies targeted at the adaptive immune response has indicated that other factors are at play [33]. Indeed, many current studies have investigated the involvement of beta cells in their own demise. We, along with other groups, have recently shown that beta cells can express inflammatory molecules and cytokines such as IL-17 [30] and HLA-II [13, 34]. We’ve also previously demonstrated that beta cell HLA-II expression can be induced *in vitro* even in healthy islets with combinations of proinflammatory cytokines (IFN-γ, TNF-α, IL-1β) that macrophages are known to secrete [13]. Interestingly, we’ve found that macrophages in patients with type 1 diabetes highly express HLA-II, suggesting that they are activated, and this is a predominantly islet-specific phenomena. Furthermore, macrophages highly expressing HLA-II are less abundant in the non-diabetic or even Aab+ donors. Although Aab+ donors had a higher proportion of CD68+ macrophages when compared with the non-diabetic donors, they did not have a higher proportion of macrophages highly expressing HLA-II. There seems to be an accumulation of macrophages in the pancreas at the early stages of disease progression, but we are unable to definitively determine at what point HLA-II becomes upregulated and how this upregulation influences macrophage function and their subequent effect on beta cell decline. Although we are limited by the markers we used from determining with certainty whether this subset of HLA-II upregulated islet macrophages is performing an innate effector or tolerogenic function, murine macrophages which express a high intensity of MHC-II have been shown to induce a Th1 response which suggests that they are more likely to be of an M1 phenotype [35]. Islet macrophages in the NOD model, even in steady state conditions, exhibit a high expression of MHC-II, IL-1β and TNF-α transcripts, which is characteristic of an M1 activation phenotype [36]. Additionally, single-cell RNA sequencing analysis in the NOD model has identified a subset of pro-inflammatory macrophages (Cxcl9 expressing macrophages), that are highly pathogenic in diabetes initiation. These macrophages increased rapidly between the period of early T cell infiltration and clinical onset of autoimmune diabetes [37], and showed an increased expression of pro-inflammatory factors such as IL-12 and IFN-γ inducible genes, costimulatory molecules, and MHC-I and MHC-II molecules, compared to other subsets of macrophages. What is more, adoptive transference of M2 macrophages in the NOD mice prevented diabetes onset; those macrophages homed into the pancreas promoting beta cell survival [38].

In the type 1 diabetes donors we’ve observed a strong correlation between the amount of islet & peri-islet macrophages highly expressing HLA-II and the percentage of HLA-II+ beta cells. Taken together, the increased infiltration of macrophages with proximity to the islets of patients with type 1 diabetes, their increased level of HLA-II expression–suggesting activation of an M1 phenotype–and their correlation with HLA-II+ beta cells, could imply that they are involved in disease pathogenesis and in the induction of aberrant HLA-II expression by the beta cells in type 1 diabetes. However, it’s important to note that macrophages represent about 50% of islet resident immune cells in humans, and so other leukocytes, such as dendritic cells, could also contribute to this dysfunction [39].

Pancreatic macrophage infiltration is characteristic of type 1 diabetes and their accumulation begins even in the pre-diabetic stages suggesting their involvement in disease initiation. The events that lead to this macrophage accumulation are still to be determined. Our evidence indicates that an increased presence of macrophages highly expressing HLA-II in the islet and islet periphery of type 1 diabetes donors are characteristic of dysregulation of innate immune interactions with the islet, and our hypothesis is that these cells are also involved in type 1 diabetes progression. The mechanism and factors that skew islet macrophage activation in type 1 diabetes, and at what point in disease progression this occurs have yet to be elucidated. Future research into these issues may provide useful information to design effective therapeutic targets to attenuate macrophage infiltration and activation in the pancreas which could possibly extend or preserve local immune tolerance.

## Acknowledgements

The authors wish to thank nPOD for their collaboration in their study. We would like to thank all organ donors and their families, whose generosity made this study possible. This research was performed with the support of the Network for Pancreatic Organ donors with Diabetes (nPOD; RRID:SCR_014641), a collaborative type 1 diabetes research project sponsored by JDRF (nPOD: 5-SRA-2018-557-Q-R) and The Leona M. & Harry B. Helmsley Charitable Trust (Grant #2018PG-T1D053). Organ Procurement Organizations (OPO) partnering with nPOD to provide research resources are listed at http://www.jdrfnpod.org//for-partners/npod-partners/.

## Funding

This research was supported by National Institute of Health grant R01AI134971 and R01AI092453. The Zeiss LSM 880 was funded by NIH S10 OD021831. EQ-M. was supported by a postdoctoral grant of Fundación Martín Escudero. S.M. was funded by an Imaging Scientist grant (2019-198153) from the Chan Zuckerberg Initiative.

## Author Contributions

EQ-M conceived and designed the study, performed experiments, designed the image analysis strategy, analyzed and interpreted data, and wrote the manuscript. SZ, TC, and SR performed experiments, analyzed data, wrote and critically revised the manuscript. SM wrote groovy scripts, participated in designing image analysis strategy, and critically revised the manuscript. WK and ZM participated in designing image analysis strategy and critically revised the manuscript. MA participated in the design of the study and critically revised the manuscript. MvH designed the study, interpreted data, and revised the manuscript. MvH and EQM are guarantor of this work and, as such, had full access to all the data in the study and take responsibility for the integrity of the data and the accuracy of the data analysis. All the authors approved the final version of the manuscript to be published.

## Materials Availability, and Data and Code Availability

- This study did not generate new unique materials or reagents.
- All data reported in this paper will be shared by the lead contact upon request.
- All original code has been deposited by our collaborator Sara McArdle at https://github.com/saramcardle and is publicly available as of the date of publication.
- Any additional information required to reanalyse the data reported in this paper is available from the lead contact upon request.

## Declaration of Interests

MvH is employed as Vice President and Senior Medical Officer at Novo Nordisk and he is currently employed at the University of Miami. SZ is currently an employee of Gilead Sciences. The authors declare no other conflicts of interest in relation with this study.

