## Supplementary Materials for "Whole tissue spatial cellular analysis reveals increased macrophage infiltration in pancreata of autoantibody positive donors and patients with type 1 diabetes"

Supp. Table 1: Clinical Characteristics of Pancreatic Organ Donors

| Disease | nPOD # | Age (years) | BMI ($kg/m^2$) | Sex | Time in ICU (days) | C-peptide (mmol/L) | T1D duration (years) | Autoantibodies |
|---|---|---|---|---|---|---|---|---|
| Non-diabetic | 6029 | 24 | 22.6 | F | – | – | – | – |
| Non-diabetic | 6073 | 19.2 | 36 | M | 4.47 | 0.69 | – | – |
| Non-diabetic | 6098 | 17.8 | 22.8 | M | 1.74 | 1.41 | – | – |
| Non-diabetic | 6251 | 33 | 29.5 | F | 3.73 | 1.92 | – | – |
| Non-diabetic | 6271 | 17 | 24.4 | M | 0.48 | 11.47 | – | – |
| Aab+ | 6123 | 23.2 | 17.6 | F | 3.55 | 2.01 | – | GADA |
| Aab+ | 6147 | 23.8 | 32.9 | F | 2.65 | 3.19 | – | GADA |
| Aab+ | 6167 | 37 | 26.3 | M | 2.95 | 5.43 | – | IA2A, ZnT8A |
| Aab+ | 6197 | 22 | 28.2 | M | 2.96 | 17.48 | – | GADA, IA2A |
| Aab+ | 6301 | 26 | 32.08 | M | 3.16 | 3.92 | – | GADA |
| T1D | 6039 | 28.7 | 23.4 | F | – | <0.05 | 12 | GADA, IA2A, mIAA, ZnT8A |
| T1D | 6084 | 14.2 | 26.3 | M | 2.48 | <0.05 | 4 | mIAA |
| T1D | 6198 | 22 | 23.1 | F | 21.71 | <0.05 | 3 | GADA, IA2A, mIAA, ZnT8A |
| T1D | 6380 | 11.6 | 14.6 | F | 4.02 | 0.22 | 0 | – |
| T1D | 6399 | 17.42 | 32 | M | – | 1.41 | 0 | GADA, IA2A, ZnT8A |

Aab+, autoantibody-positive; F, female; GADA, GAD autoantibodies; IA2A, insulinoma-associated protein 2 autoantibodies; M, male; mIAA, micro-insulin autoantibodies; nPOD #, nPOD identification number; T1D, type 1 diabetes; ZnT8A, zinc transporter 8 autoantibodies.

Supp. Fig. 1: Visual representation of regional and spatial cell distribution analysis methodology

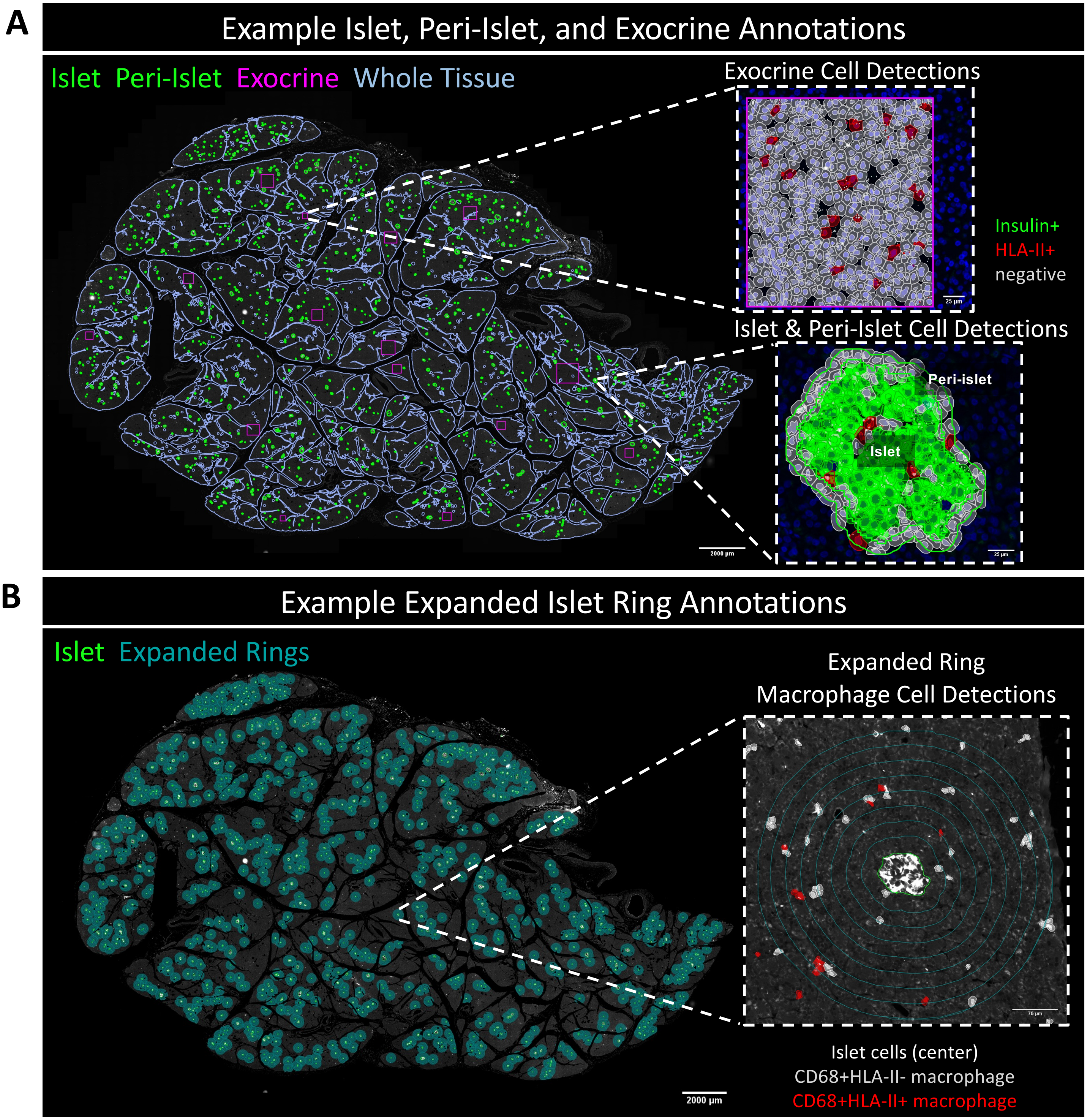


The spatial distribution of macrophages within each pancreatic tissue section was analyzed with two different semi-automated methods using machine learning algorithms and custom groovy scripts in QuPath. **(A)** Whole tissue annotation (purple). Regional analysis with automatically detected islet (green) and peri-islet (green; 10 µm radius) regions and selected exocrine regions (pink). Both amplified images show the cell classification in regions of interest. **(B)** Islet proximity analysis with automatically detected islet regions (green) and expanded islet regions consisting of 8 concentric rings (teal; 25 µm radius). For both methods, the number, percentage and density of cells were counted within each region and categorized by their classification (positive or negative) for each marker (insulin, CD68, and HLA class II).

Supp. Fig. 2: Macrophage infiltration is greatest in the pancreas of donors with type 1 diabetes regardless of region

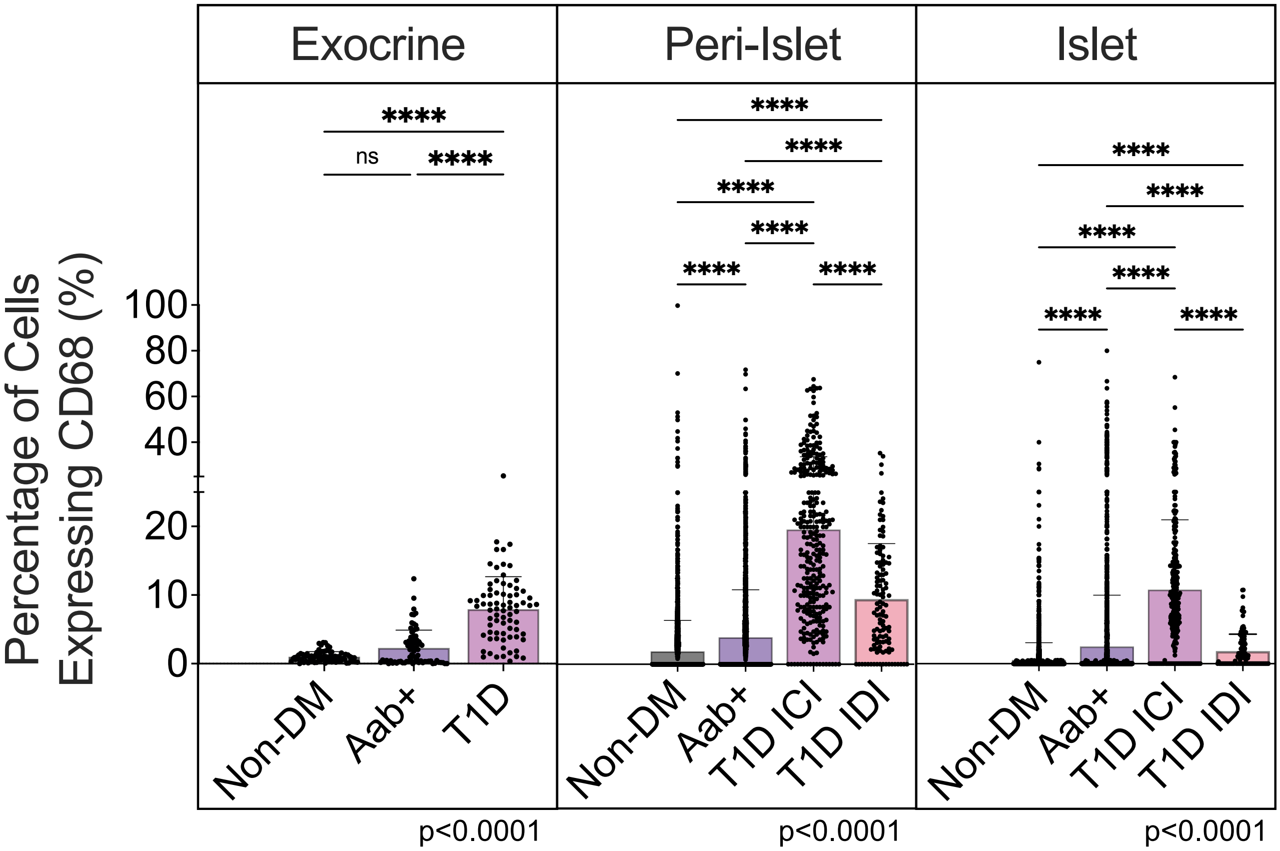


The percentage of total cells which express CD68 is higher in the insulin-containing islets (ICI) and associated peri-islet (10 µm radius) and exocrine regions of the T1D cases than any other group. The differences observed between groups were statistically significant for the exocrine, peri-islet and islet regions ($p<0.0001$). Each datapoint represents one region (i.e., one islet), and the mean + SD is shown. Statistical significance was calculated using Kruskal-Wallis test, and multiple comparisons were assessed using Dunn's test (**** $p<0.0001$, ns not-significant).

Supp. Fig. 3: Macrophage distribution near islets varies within and between groups

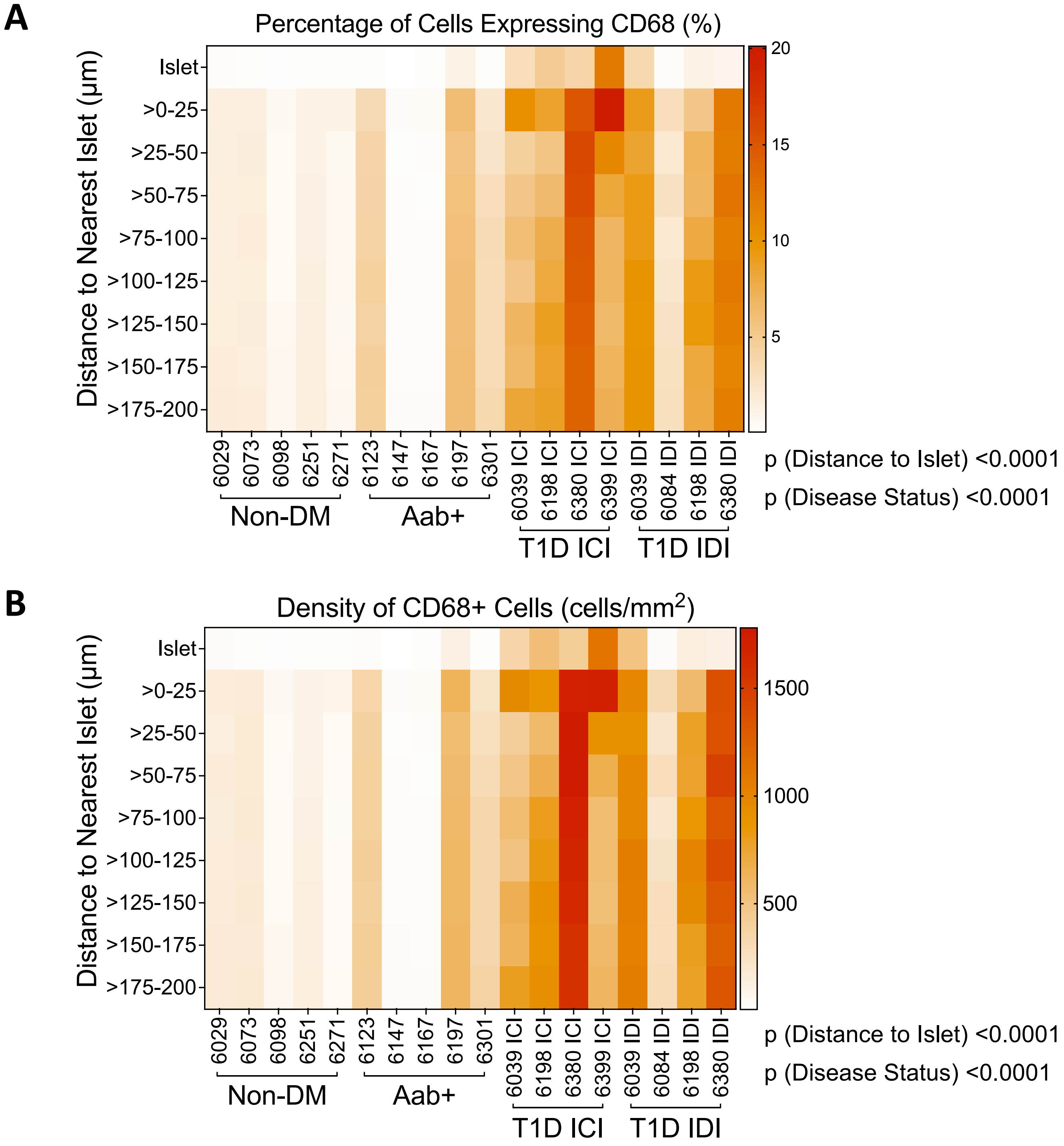


Case-wise representation of the expanded islet region analysis consisting of 8 concentric rings (25 μm radius). Both the **(A)** percentage and **(B)** density of cells expressing CD68 within each ring varied between cases and groups. Statistical significance was calculated using Two-way ANOVA test, and multiple comparisons were assessed using either Tukey's test.

Supp. Fig. 4: Distance-wise variation of macrophage distribution for expanded ring spatial analysis

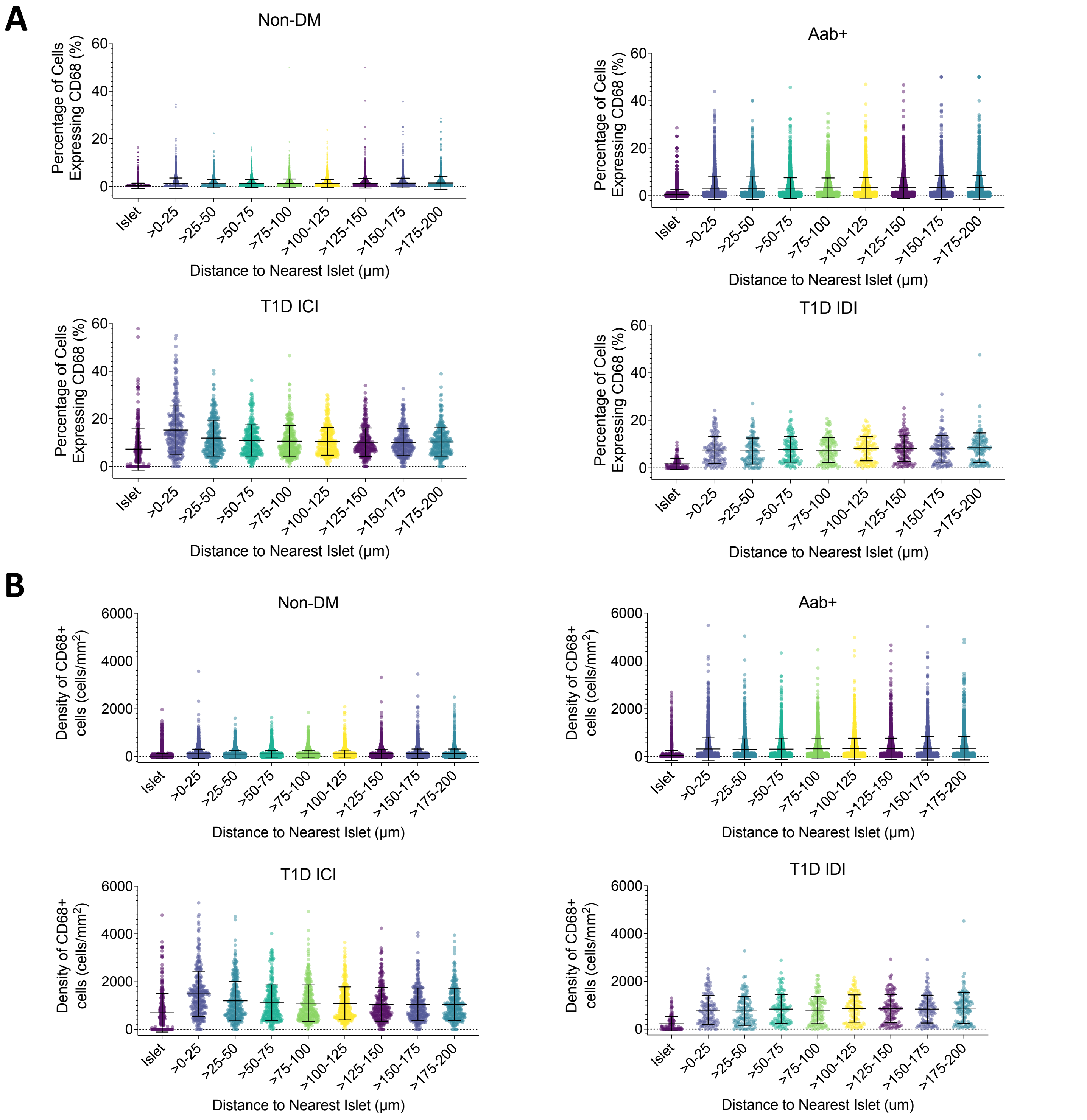


Raw data of the expanded islet region analysis consisting of 8 concentric rings (25 μm radius) for the **(A)** percentage and **(B)** density of cells expressing CD68 within each ring. Each datapoint represents one region and the mean ± SD is shown.
